# Theta-burst transcranial magnetic stimulation to the prefrontal or parietal cortex does not impair metacognitive visual awareness

**DOI:** 10.1101/058032

**Authors:** Daniel Bor, David J. Schwartzman, Adam B. Barrett, Anil K. Seth

## Abstract

Neuroimaging studies commonly associate dorsolateral prefrontal cortex (DLPFC) and posterior parietal cortex with conscious perception. However, such studies only investigate correlation, rather than causation. In addition, many studies conflate objective performance with subjective awareness. In an influential recent paper, Rounis and colleagues addressed these issues by showing that theta burst transcranial magnetic stimulation (tbs-TMS) applied to the DLPFC impaired metacognitive (subjective) awareness for a perceptual task, while objective performance was kept constant. We attempted to replicate this finding, with minor modifications, including an active tbs-TMS control site. Using a between-subjects design for both DLPFC and posterior parietal cortices, we found no evidence of a tbs-TMS-induced metacognitive impairment. In a second experiment, we devised a highly rigorous within-subjects tbs-TMS design for DLPFC, but again failed to find any evidence of metacognitive impairment. One crucial difference between our results and the Rounis study is our strict exclusion of data deemed unsuitable for a signal detection theory analysis. Indeed, when we included this unstable data, a significant, though invalid, metacognitive impairment was found. These results cast doubt on previous findings relating metacognitive awareness to DLPFC, and inform the current debate concerning whether or not prefrontal regions are preferentially implicated in conscious perception.

## Introduction

Many studies support the view that the lateral prefrontal cortex, as well as the posterior parietal cortex (PPC), are associated with conscious processes [1-11](See Bor & Seth, 2012; Dehaene & Changeux, 2011; Koch et al, 2016 for reviews). However, the vast majority of these studies, employing neuroimaging techniques, are correlational, and therefore are unable to test whether the prefrontal parietal network is causally implicated in conscious perception. Prefrontal and parietal lesion studies could in contrast demonstrate a causal relationship between this cortical network and consciousness. However, such studies have produced more equivocal results, and tend to show at best subtle impairments in conscious detection [12,13]. It is possible, however, that these cortical regions show especially plastic responses to damage, thus protecting individuals from cognitive and conscious impairments [14].

Transcranial Magnetic Stimulation (TMS) provides an alternative method for investigating whether a specific brain region is necessary for a certain function, by temporarily disrupting localised neuronal activity for seconds or minutes. An advantage of TMS, besides its non-invasive nature, is that TMS-induced changes are limited to short time periods so that more long-term, uncontrolled-for, plastic changes that are possible in lesion studies are not an issue. Studies using this technique applied to the dorsolateral prefrontal cortex (DLPFC) [15] and right PPC [16] have demonstrated impairments in conscious change detection. In addition, TMS to the PPC has been shown to decrease switch rate in a binocular rivalry paradigm [17]. However, studies like these tend to use repetitive TMS, whose peripheral consequences (e.g. noise, facial nerve stimulation) could itself create distractions that cause transitory cognitive impairments for the ongoing task. Furthermore, it is commonly difficult in such studies to disentangle conscious effects from lower level changes: for instance, impairments in change detection could arise if TMS disrupted the unconscious processing of basic visual features.

Rounis and colleagues [18] designed a study to overcome these issues. First, continuous theta burst TMS (cTBS) was used instead of conventional repetitive TMS. This technique involves a very rapid sequence of TMS pulses, typically for 40 s. The protocol used in our study is thought to suppress cortical excitability for up to 20 minutes [19]. In this way, TMS administration can be entirely separated from the behavioural task, and therefore will not distract the participants from it. Second, Rounis and colleagues used a metacontrast mask binary perceptual task with stimulus contrast titration in order to maintain objective performance at 75% accuracy. By combining this design with advanced methods in signal detection theory (SDT) [20-22], they were able to isolate the effects of TMS-induced inhibition of DLPFC on metacognitive sensitivity.

Metacognition tracks the extent to which an individual is aware of their own knowledge, commonly in mnemonic or perceptual domains, by assessing how closely confidence relates to decision accuracy. Since metacognitive sensitivity, in humans at least, is typically assumed to index the extent of subjective awareness (of one’s own mental states), the Rounis study used a particularly rigorous method to explore changes in conscious perception resulting from transient deactivation of specific cortical regions. Neuroimaging and electrophysiological studies have previously linked either lateral prefrontal cortex [23-26] or posterior parietal cortex [27] with metacognitive processes. In addition, a small (n=7) patient lesion study showed that the anterior prefrontal cortex (i.e. a region neighbouring the DLPFC) selectively impaired perceptual metacognition, though not memory-based metacognition, compared with patients who had temporal lobe lesions [28]. However, Rounis and colleagues were the first to provide persuasive non-patient-based evidence that DLPFC has a key causal role to play in reportable conscious perception, by showing that cTBS to DLPFC, but not sham cTBS, reduced metacognitive sensitivity for the perceptual task, while objective sensitivity remained unchanged.

Given that the Rounis study is one of the most definitive to have indicated a causal link between DLPFC and metacognitive sensitivity, it is somewhat surprising that it has not yet been replicated. In experiment 1 we therefore sought to replicate the Rounis study, as well as extend it to the posterior parietal cortex, since this region in neuroimaging studies is very commonly co-activated with DLPFC, both in studies of conscious perception [2,3] and more widely for many cognitive processes [1,29-33]. In addition, we included extra conditions where TMS was either only applied to the left or right hemisphere, so that we could explore laterality effects. Furthermore, we attempted to enhance the original Rounis design, by including an active TMS control (vertex), rather than sham stimulation. In experiment 2, we attempted for a second time to replicate the Rounis study, copying their design more closely using a within subjects design, by examining cTBS to bilateral DLPFC, as they did, though still with an active control instead of sham.

## Experiment 1

This experiment was a direct replication and extension of the Rounis paradigm [18], except that a between subjects design was used. Each volunteer was assigned to one of 5 TBS groups: i) bilateral DLPFC, ii) bilateral PPC, iii) left DLPFC and PPC, iv) right DLPFC and PPC, and v) VERTEX (control). Other minor deviations from the previous protocol are described below. All such minor deviations were carefully considered to improve the chances of detecting valid effects, as we explain in each case.

## Methods

### Participants

90 healthy volunteers (49 women, mean age 22.7, SD age 5.1), with no history of neurological disorders, psychiatric disorders, or head injury were recruited from the local student population. Written informed consent was obtained from all volunteers. The study was approved by the University of Sussex local research ethics committee. Methods were carried out in accordance with the approved guidelines.

### Experimental design

The experimental design was taken directly from Rounis and colleagues [18], who also generously provided the experimental software, which was a COGENT program, running under Matlab. Participants performed a two-alternative forced choice task (Figure 1A). All testing was carried out in a darkened room. Stimuli were presented approximately 40 cm distance from the volunteers’ eyes on a CRT monitor with a 120 Hz refresh rate. Black stimuli were presented on a white background. During each trial, a square and a diamond 0.8 degree wide each were presented for 33ms 1 degree either side of a central fixation cross. 100 ms after stimulus onset, a metacontrast mask was presented for 50ms. Participants had to identify whether the diamond had appeared on the left and square on the right, or vice versa (this is the perception, or type I task). Both stimulus possibilities were presented in pseudorandom order with equal probability. Simultaneously, volunteers provided subjective stimulus ratings (this is the metacognitive, or type II task). In the Rounis paradigm [18], participants were asked to make a relative distinction between “clear” and “unclear” ratings, in the context of the experiment as a whole. This was designed to generate roughly equal answers for each rating, so as to make the SDT analyses more stable (personal communication). However, from summary data kindly supplied by Rounis and colleagues, 13/20 participants in their study had at least one session with unstable data (where type I or II false alarm rate (FAR) or hit rate (HR) were >0.95 or <0.05). Therefore, based on our exclusion criteria, we only would have included 7/20 of their subjects for analysis, indicating that their strategy for ensuring stable data were, by our criteria, not successful.

**Figure 1.**
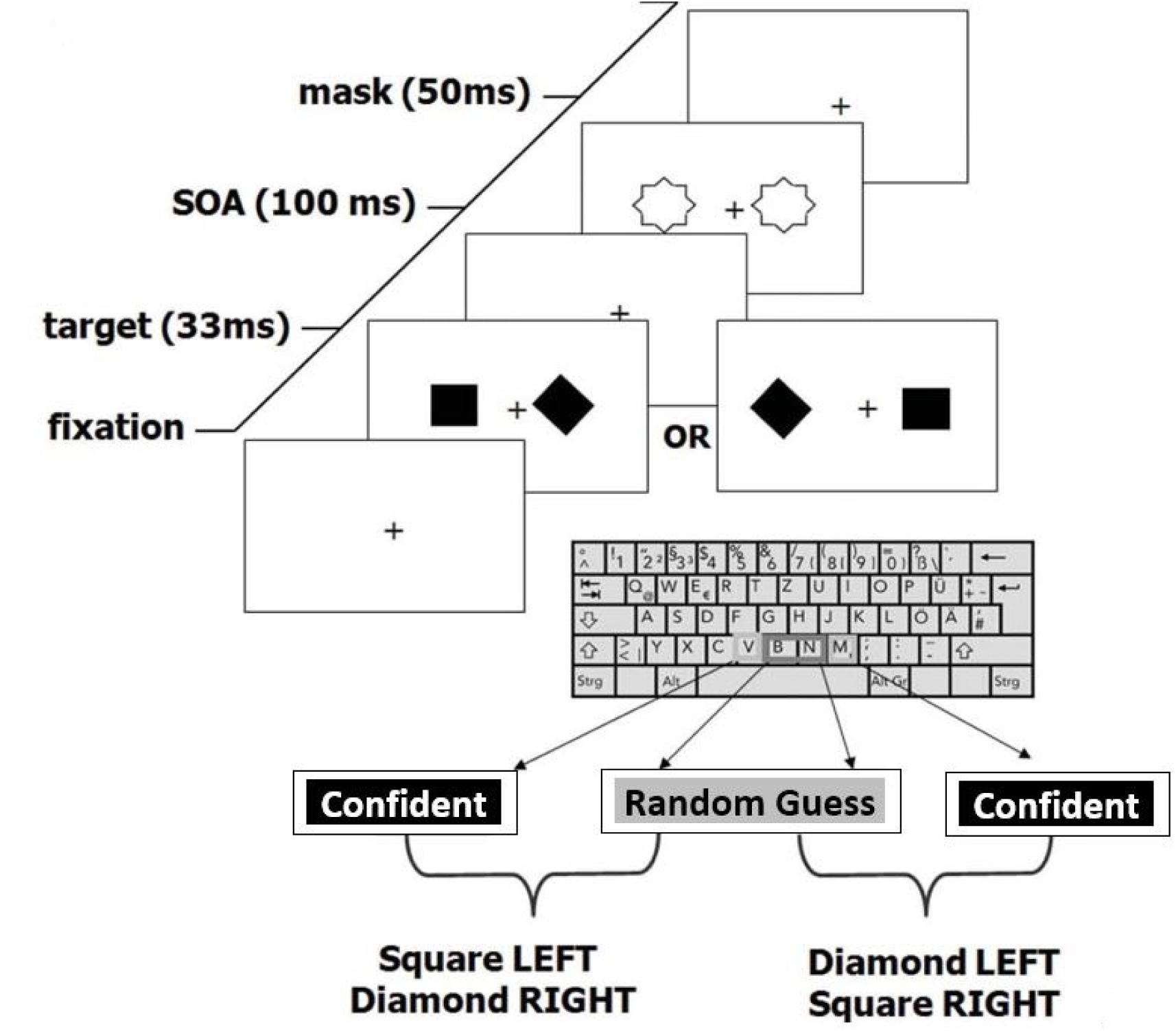
Experimental design was identical to Rounis and colleagues [18], apart from exceptions described in methods. Most notably, confidence in choice was used instead of visibility to determine metacognitive judgement. Participants were presented with either a diamond on the left and square on the right or vice versa, followed by a metacontrast mask. They were then required to make a combined judgement as to the stimulus configuration and their level of confidence in that decision. Adapted from Rounis [18] with permission.

We were concerned that managing the relative frequency of subjective ratings of “clear” and “unclear” labels across an experiment may have placed additional working memory demands on participants, since they would need to keep a rough recent tally of each rating in order to balance them out. In addition, these labels were difficult to interpret psychologically on account of their relative nature. We therefore opted instead for the labels “[completely] random [guess]” and “[at least some] confidence.” Using confidence instead of clarity labels is a common practice, consistent with other recent metacognition studies [25,26]. Although this method could potentially introduce more unstable SDT values into the analysis, given that participants could in principle give all answers as “random” or “confidence,” we excluded this possibility by removing from the analysis any volunteers who had any HR and FAR values below 0.05 or above 0.95 (i.e. beyond the cut-off points for obtaining stable z-transforms from which to compute SDT quantities; see Discussion and Barrett et al [2013]).

Each subject attended a single testing session, which began with an easy demonstration phase of 100 trials, followed by a practice phase, also of 100 trials. The practice phase was designed to further familiarize participants to the experiment, and allow them to reach a steady state of performance. Objective performance was controlled to be close to 75% throughout the experiment, by titrating the contrast levels of the stimuli (with black the easiest contrast and a very light grey the hardest, all against a white background) using a staircase procedure [21]. Each trial was randomly assigned to either staircase A or B. For staircase A, current trial contrast was increased (i.e. darkened) if the participant responded incorrectly on the previous staircase A trial, and contrast was decreased (i.e. made lighter), if the volunteer had correctly responded on the previous two staircase A trials. Staircase B worked in the same way, except that three prior consecutive correct responses were required to reduce contrast. Contrast changes were made in 5% increments.

We were concerned that the Cogent experimental script of Rounis and colleagues [18] could, under certain circumstances, fail to allow participants to reach a steady state during the practice phase. Therefore we made minor changes to the script and paradigm at this stage: we removed a small bug in the script, which caused the contrast levels to jump erratically at contrast levels close to the most difficult end; we changed the practice staircase procedure to be identical to that of the main experimental blocks (previously it was significantly easier than the main blocks, potentially leading to the steady state of participants set wrongly for the main blocks); we also, unlike the Rounis paradigm, occasionally repeated the practice block if it was clear from the performance graphs that the participant hadn’t yet reached a steady state in performance; finally we noted in piloting the experiment that a small group of participants were at ceiling on the task. Therefore we extended the contrast range: when participants were at the 95% white level (previously the final contrast setting), further 1% contrast increments were introduced, up to 99% white.

After the practice stage, volunteers carried out a pre-TBS block of 300 trials, to measure baseline subjectivity ratings. Brief breaks were allowed after every 100 trials. The block took approximately 11 minutes to complete. After the pre-TBS block was completed, the TBS pulses were administered. Following TBS administration, a further post-TBS block of 300 trials was administered.

In addition, the Cattell Culture Fair IQ test 2a was given to participants before the main experiment, in order to explore the modulatory effects of IQ on metacognition. Note that due to time constraints, approximately a quarter of participants were unable to take the IQ test.

### Theta-burst stimulation

A Magstim Super Rapid Stimulator (Whitland, UK), connected to four booster modules with a standard figure of eight coil, was used to administer the TBS TMS pulses. For each TBS session, a stimulation intensity of 80% of active motor threshold (AMT) for the left dorsal interosseous hand muscle was used. The AMT was defined as the lowest intensity that elicited at least 3 consecutive twitches, stimulated over the motor hot spot, while the participant was maintaining a voluntary contralateral finger-thumb contraction. cTBS was delivered with the handle pointing posteriorly and the coil placed tangentially to the scalp. The standard cTBS pattern used, as with the Rounis study, was a burst of three pulses at 50 Hz given in 200 ms intervals, repeated for 300 pulses (or 100 bursts) for 20 s. Following a 1 minute interval, this was repeated at a different site for a further 20s (or again on the vertex in the control condition), determined by which group the participant was assigned to. The five groups were: i) bilateral DLPFC, ii) bilateral PPC, iii) left DLPFC and PPC, iv) right DLPFC and PPC, and v) VERTEX (control). Previous studies have demonstrated that this cTBS procedure, when applied to the primary motor cortex, induces a decrease in corticospinal excitability lasting about 20 minutes [19]. Where stimulation involved two sites (all except the VERTEX group), the choice of first stimulation site was counterbalanced between participants.

The DLPFC site was located, as with the Rounis study, 5cm anterior to the “motor hot spot”, on a line parallel to the midsagittal line. The PPC site was located in the same way as the DLPFC site, except for being 5cm posterior to the “motor hot spot.” The “motor hot spot” was defined functionally as the maximal evoked motor response, when determining AMT.

### Data Analysis

Following Rounis and colleagues [18], a range of measures were used to assess the change in metacognitive performance between the pre- and post- TMS blocks. This included the phi correlation between accuracy and subjective ratings, as well as meta d’, an SDT measure thought to reflect the amount of signal available for a participant’s metacognitive disposal. There are specific methodological advantages provided by meta d’, as compared to type II d’, for measuring metacognitive sensitivity [18,20,22]. In particular, it is well known that type II d’ is highly dependent on both type I and type II response bias whereas meta d’ is approximately invariant with respect to changes in these thresholds and thus provides a more direct measure of metacognitive sensitivity [20,22]. For further discussion and detailed computational analysis of different methods to measure metacognitive sensitivity, see Barrett et al (2013) and [34].

We followed the Rounis approach to generate two estimates of meta d’, based on the participant’s type II HR and FAR, conditional on each stimulus classification type. The two estimates were combined using a weighted average, based on the number of trials used to calculate each estimate. There are currently two approaches to generate meta d’ values: sum of squared errors (SSE) and maximum likelihood estimates (MLE). Here we report SSE, as in the Rounis paper, although MLE results were also analysed and yielded very similar values. For completeness, we also report type II d’ results, although we recognise that this measure has methodological disadvantages compared with meta d’ [20,22].

In summary, using correlational, type II d’ and meta d’ approaches, we tested for any reduction in metacognitive sensitivity following administration of cTBS. The comparison of the DLPFC group with the vertex control group on this measure was a direct attempt at replicating the Rounis paradigm [18], although in our case a between groups design and an active, rather than sham, control was used. Following Rounis, we report 1-tailed values, due to directional hypotheses that metacognitive sensitivity will be reduced following TBS to any non-control pair of sites.

## Results

Although in the Rounis paradigm no participants were excluded, in our study, for each group, subjects were excluded from the analysis if: i) in either of the two sessions there were extreme SDT values for type I or 2 HR and FAR (<0.05 or >0.95); ii) in either of the two sessions accuracy was significantly below the 75% required (at least 10% lower): or iii) because of problems with the TMS administration, for instance that the experimenter was unable to find an accurate AMT. See Table 1 for a summary. Note that our proportion of subjects having extreme SDT values was considerably less than in the Rounis study (27/90 (30%) compared to 13/20 in the Rounis study (65%)), though given in their within-subjects design participants had 4 TMS sessions instead of 2, our results, in terms of proportion of subjects excluded per TMS session, are roughly comparable to theirs.

**Table 1.** List of inclusions and exclusions for experiment 1 participants

| **Group** | **Original n** | **Excluded due to extreme (unstable) SDT values** | **Excluded due to Accuracy problems** | **Excluded due to TMS administration problems** | **Remaining n for analysis** |
| --- | --- | --- | --- | --- | --- |
| **Bilateral DLPFC** | 17 | 4 | 0 | 1 | 12 |
| **Bilateral PPC** | 16 | 3 | 1 | 2 | 10 |
| **Left DLPFC and PPC** | 18 | 6 | 1 | 1 | 10 |
| **Right DLPFC and PPC** | 21 | 9 | 1 | 2 | 9 |
| **Vertex (control)** | 18 | 5 | 0 | 1 | 12 |
| **TOTALS** | 90 | 27 | 3 | 7 | 53 |

Unsurprisingly, given that accuracy was dynamically controlled throughout the experiment, to approximate to 75% correct, there was no difference between accuracy levels (Figure 2a) before or after TMS (F(1,48)=0.67, p>0.1; effect size: partial eta^2^ = 0.013), nor was there a TMS stage x group interaction for performance (F(4,48)=0.26, p>0.1; effect size: partial eta^2^ = 0.021).

**Figure 2.**
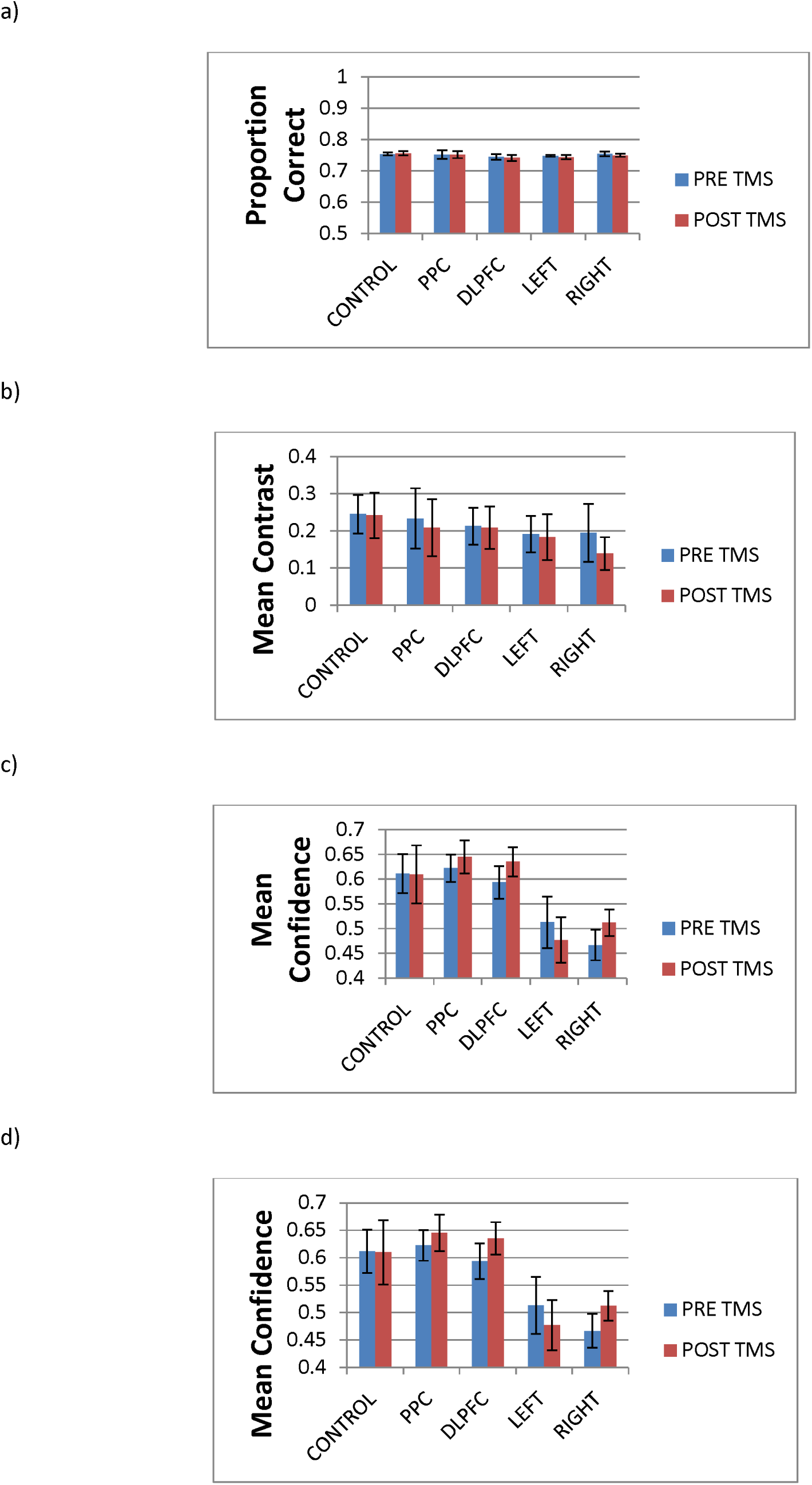
Pre and post-TMS performance measures for the different groups. a) Proportion correct. B) Mean contrast C) Mean confidence D) Reaction Time for correct responses. DLPFC = bilateral DLPFC group, PPN = bilateral posterior parietal cortex group, LEFT = left posterior parietal cortex and DLPFC group, RIGHT = right posterior parietal cortex and DLPFC group. All error bars are SE.

A more interesting comparison is the mean contrast level to keep accuracy constant at 75%. In the Rounis study [18], the mean contrast level was, on average, more difficult (lower) in the post-TMS stage, compared with the pre-TMS stage, for both real and sham TMS. In the present study, in contrast, we found no reduction in mean contrast levels (Figure 2b) between stages (F(1,48)=2.46, p>0.1; effect size: partial eta^2^ = 0.049), nor a TMS stage x group interaction for contrast levels (F(4,48)=0.61, p>0.1; effect size: partial eta^2^ = 0.048). Similarly, the Rounis study reported a decline in the fraction of stimuli that were visible (analogous to the confidence ratings in this study) following TMS treatment (independent of whether it was real or sham), but in the current study, we found neither a decline in confidence (Figure 2c) following TMS (F(1,48)=1.36, p>0.1; effect size: partial eta^2^ = 0.028), nor a TMS stage x group interaction for confidence (F(4,48)=1.44, p>0.1; effect size: partial eta^2^ = 0.107). Rounis and colleagues attributed the changes they observed to a possible “learning effect”, although another possible factor may have been an overly easy practice stage, which would have led to the main blocks having too liberal a starting contrast level. This in turn would have decreased the likelihood of stable contrast levels, especially in the first (pre-TMS) block. Therefore at least part of the reason for their “learning effect” could have been that a portion of the first block involved a transition to a stable contrast. In any case, our data show that our modifications to ensure stable contrast values at the start of the main block were effective.

In contrast to these differences, consistent with the Rounis study [18] we found faster RTs for correct responses (Figure 2D) in the second session (F(1,48)=9.36, p=0.004; effect size: partial eta^2^ = 0.163), but no interaction between RT mean session score and TMS group (F(4,48)= 1.73, p>0.1; effect size: partial eta^2^ = 0.126).

For the critical analysis of whether TMS reduced metacognitive sensitivity, we found no evidence for this in any of our groups. There was no TMS group x time interaction for the correlation between accuracy and confidence, phi (F(4,48)= 0.14, p>0.1; effect size: partial eta^2^ = 0.002), nor for meta d’ – d’ (F(4,48)= 0.06, p>0.1; effect size: partial eta^2^ = 0.005), nor type II d’ (F(4,48)= 0.162, p>0.1; effect size: partial eta^2^ = 0.013). In order to further verify this failure to replicate the Rounis results, we carried out t tests and Bayes factor analyses on the above 3 measures for all test groups against the vertex control. The Bayes factor analyses used the correlation and meta d’ priors from the Rounis study to constrain the calculation (assuming that the other experimental groups would show the same difference as the bilateral DLPFC group). These priors were -0.4 and -0.05 for the post- minus pre-TMS difference of DLPFC condition versus sham control for meta d’ – d and the accuracy-visibility correlations, respectively. However for type II d’, which wasn’t reported in the Rounis study, lower and upper bounds of the average type II d’ scores for all current sessions (0.93) were used in lieu of a prior. Another method of calculating the Bayes factor here is to take a ratio of the average type II d’ scores for both sessions to the average meta d’ – d’ scores (0.930/0.147=6.3) and multiply this by the meta d’ – d’ Rounis prior (6.3*-0.4 = -2.528), as an estimate of the type II d’ difference Rounis and colleagues would have observed. If we use this method instead, all Bayes factor scores are less than 0.2 (i.e. robust null results).

As shown in tables 2, 3 and 4, no comparison between control or experimental group approached significance, even using 1-tailed statistics, on meta d’, type II d’, and the correlation between accuracy and confidence, respectively (see also Figure 3). In addition, effect sizes were extremely small, supporting the suggestion that there were no differences between the experimental and control groups. Furthermore, the Bayes factor analyses were either approaching or lower than the lower bound of 0.33, which is considered substantial support for the null hypothesis [35]. Given the relatively small sample sizes of approximately 11 per group, the fact that the Bayes factor scores didn’t reach a robust null in some cases might be due to lack of power. This situation is partially rectified in the second experiment where data from both experiments can be combined.

**Figure 3.**
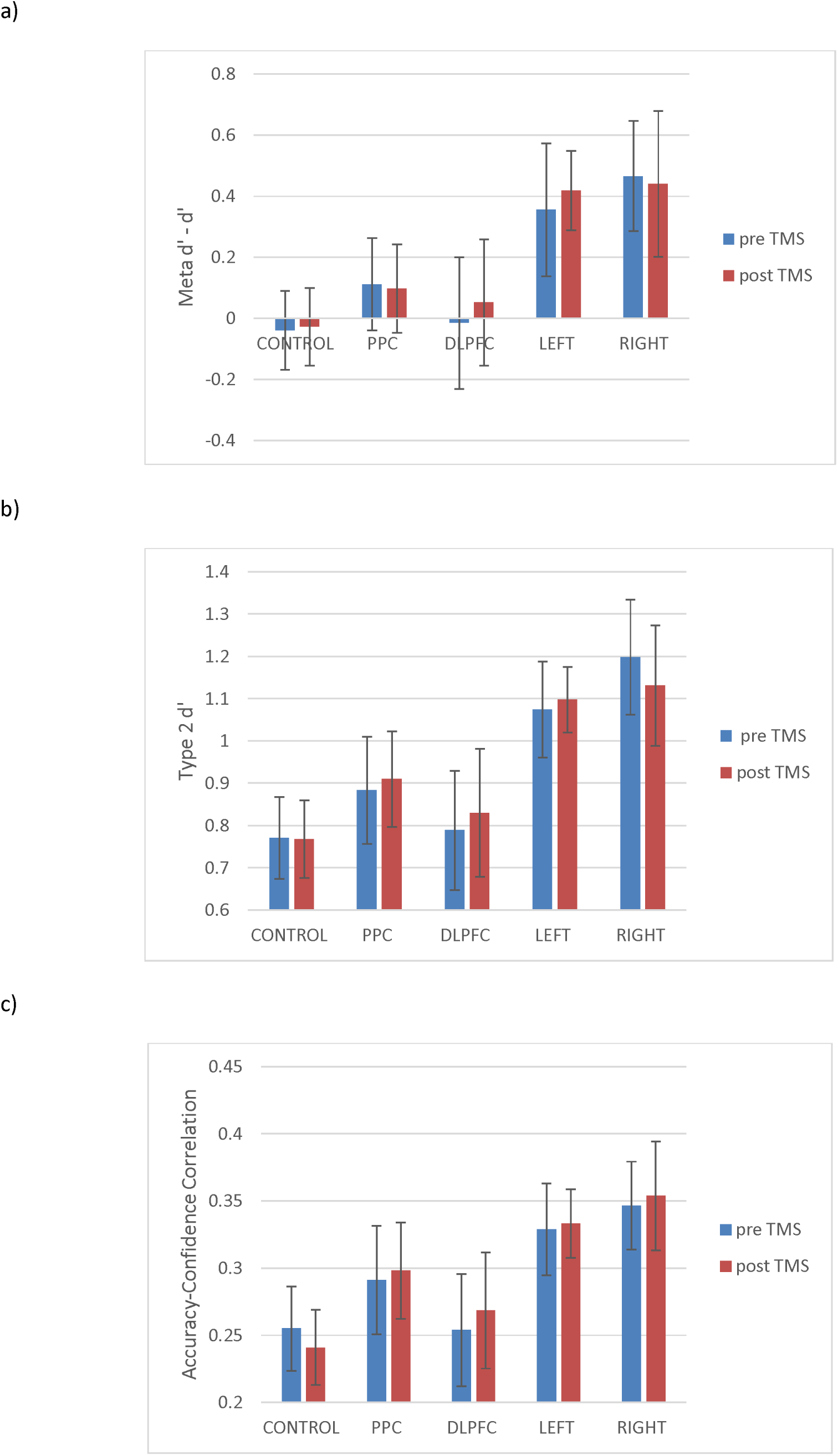
Pre- and post-TMS metacognitive measures for the different groups. a) meta d’ – d’. b) type II d’. c) Accuracy-confidence phi correlation. Group labels as figure 2. All error bars are SE.

**Table 2.**
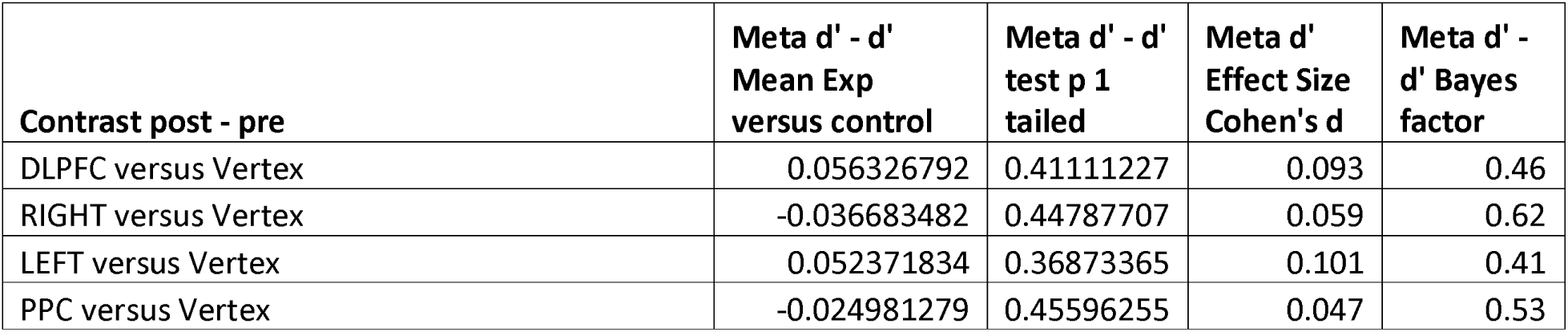
Meta d’ table of t tests, effect sizes and bayes factors analyses between conditions and control (NB for the Rounis study, post – pre TMS meta d’ -d’ Mean DLPFC versus sham control was -0.4)

**Table 3.** Type II d’ table of t tests, effect sizes and Bayes factors analyses between conditions and control (NB no type II d’ results were reported in the Rounis study. Lower/upper bounds of average type II d’ scores were used instead.)

| **Contrast post - pre** | **Type II d' Mean**<br>**Exp versus**<br>**control** | **Type II d' t**<br>**test p 1**<br>**tailed** | **Type II d'**<br>**Effect Size**<br>**Cohen's d** | **Type II d'**<br>**Bayes**<br>**factor** |
| --- | --- | --- | --- | --- |
| DLPFC versus Vertex | 0.043731728 | 0.37964982 | 0.12672943 | 0.2 |
| RIGHT versus Vertex | -0.064632552 | 0.34881974 | 0.17580987 | 0.24 |
| LEFT versus Vertex | 0.090041273 | 0.28881694 | 0.07979371 | 0.62 |
| PPC versus Vertex | 0.02878612 | 0.41544334 | 0.09177253 | 0.18 |

**Table 4.** Correlation between accuracy and confidence table of t tests, effect sizes and Bayes factors analyses between conditions and control (NB for the Rounis study, post – pre TMS accuracy-visibility correlation DLPFC versus sham control was -0.05)

| **Contrast post - pre** | **Correlation**<br>**Mean Exp**<br>**versus control** | **Correlation**<br>**(phi) t test**<br>**p 1 tailed** | **Correlation**<br>**Effect Size**<br>**Cohen's d** | **Correlation**<br>**Bayes**<br>**factor** |
| --- | --- | --- | --- | --- |
| DLPFC versus Vertex | 0.028656427 | 0.268 | 0.25704197 | 0.48 |
| RIGHT versus Vertex | 0.021489429 | 0.32 | 0.20612987 | 0.53 |
| LEFT versus Vertex | 0.018404477 | 0.31 | 0.22064111 | 0.44 |
| PPC versus Vertex | 0.021058379 | 0.31 | 0.22290338 | 0.47 |

In case there were either short-lived or delayed cTBS effects, we reran all of the above metacognitive analyses using either only the first or last 100 of the 300 trials per block. We still found no significant differences in any comparison (all p>0.2).

In order to explore the effects of including unstable values in our analysis, we generated histograms of meta d’ – d’ scores for both sessions together for all conditions, either for SDT stable data only, or for all the data (see figure 4). These suggest that adding the unstable values transforms the data from Gaussian to non-Gaussian, specifically by adding a separate group of very high meta d’ – d’ values to the sample. Using the Shapiro-Wilk test, the stable data were not significantly non-Gaussian (W = 0.988, df = 126 p=0.317). However, when considering all data, including unstable data, the Shapiro-Wilk test indicated non-Gaussianity (W = 0.871, df = 180 p<0.001), which was also true for unstable data only (W=0.887, df=54, p<0.001). Given the Rounis study did not exclude unstable subjects, there is a chance, therefore, that their data were also non-Gaussian, meaning that it would have been invalid to use parametric statistics as they did. Furthermore, we found a significant difference between the stable and unstable meta d’ – d’ values (Mann–Whitney *U* = 2552, *n*_1_ = 126 *n*_2_ = 54, *P* < 0.008 two-tailed), suggesting that data including the unstable values should not count as a single homogeneous sample.

**Figure 4.**
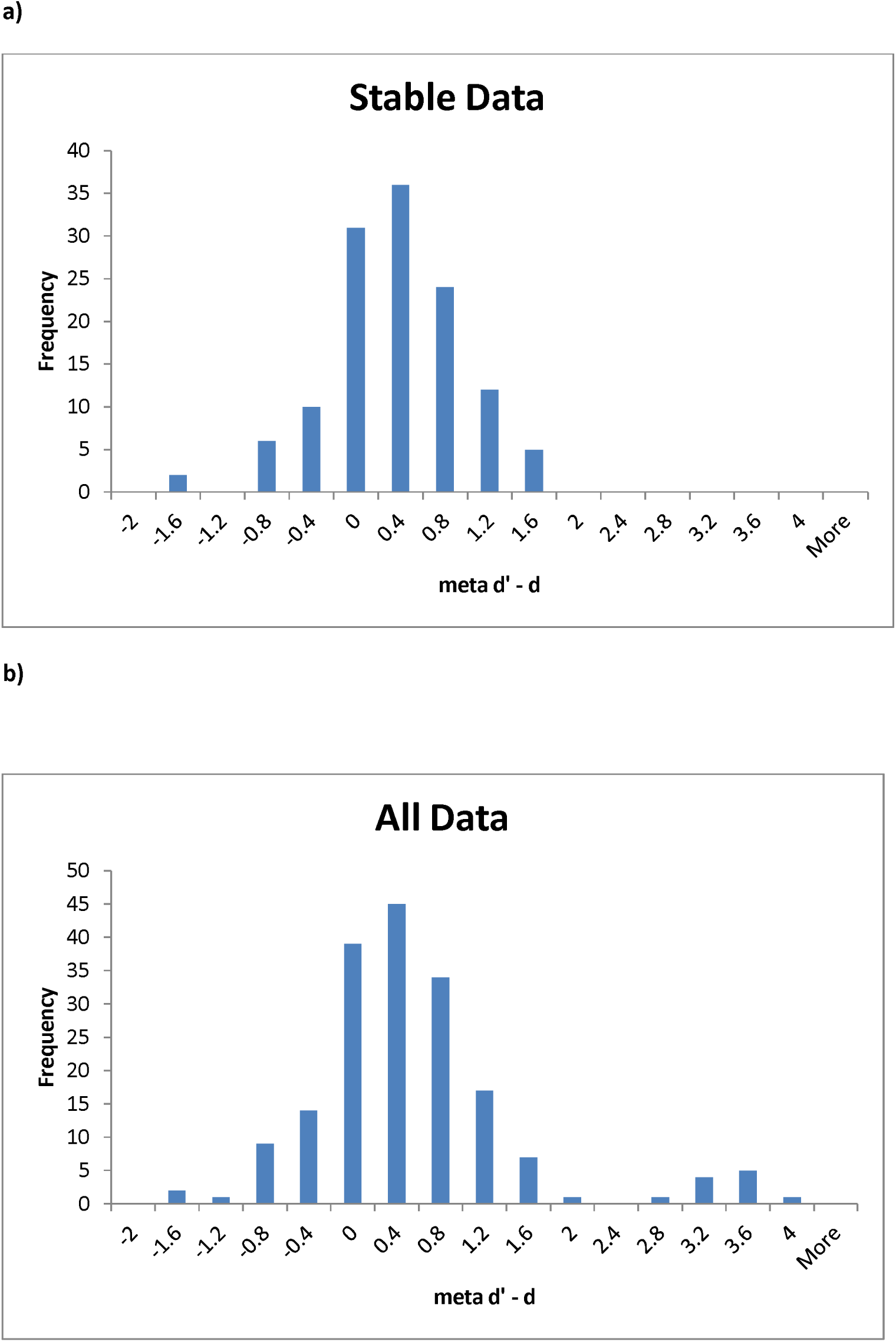
Histograms, using 0.4 sized bins, of meta d’ – d’ for a) stable data only; and b) all data (including unstable). Whereas the stable data is Gaussian, the unstable data is not.

In order to further assess potential issues with including unstable subjects, we analysed the data when including those participants we had previously excluded because of extreme HR and FAR values, using parametric statistics as in the Rounis study. Although no other effects were significant, in this analysis we did find significant differences between the DLPFC and vertex group on meta d’ – d’ scores (t(31)=1.85, p(1-tailed)=0.037; effect size Cohen’s d= 0.623). This appeared, though, to be driven more by an unpredicted boost to metacognition in the control group (0.45) than a reduction in metacognition in the DLPFC group (-0.29). We should emphasise, however, that this significant result, aside from being uncorrected for multiple comparisons, is not to be trusted as it includes data that invalidates the (parametric) assumptions underlying the analysis. We merely include this analysis to demonstrate how the inclusion of unstable values could potentially generate spurious significant results.

Finally, when exploring the relationship between IQ and metacognition, we failed to find any correlation on any of our three measures. However, we did discover a significant negative relationship between IQ and contrast level (r^2^=-0.22 t(1,40)=3.36, p=0.002; effect size Cohen f^2^=0.282) (see Figure 5), such that higher IQ participants were presented with more difficult perceptual stimuli. Similarly, there was a positive correlation between IQ and type I d’ (r^2^=-0.10 t(1,40)=2.07, p=0.045; effect size Cohen f^2^=0.111).

**Figure 5.**
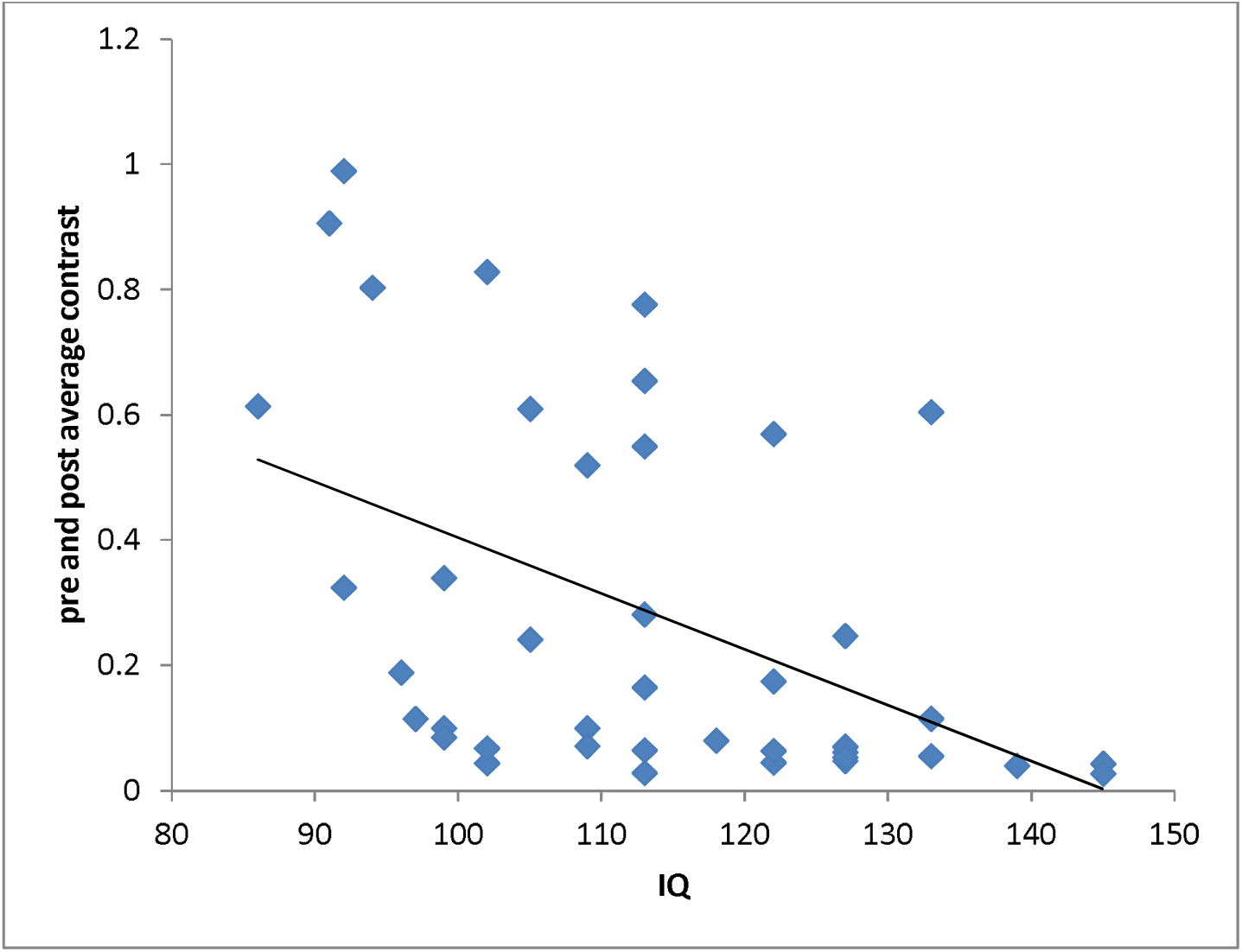
The relationship between Cattell Culture Fair IQ score and average contrast. Each blue diamond represents a single participant’s average score for both sessions. The black line is a linear best fit of the data. There was a significant negative relationship between IQ and contrast, such that higher IQ participants tended to achieve a more difficult contrast level.

## Experiment 2

Experiment 1 comprehensively failed to replicate the main result of Rounis and colleagues [18]. However, it is possible that this experiment was underpowered compared to that of Rounis and colleagues: after subject exclusions, our sample size, although more than double that used by Rounis et al, was smaller per group; in addition, we used a between subjects design, unlike the within subjects design of Rounis and colleagues. Therefore, we carried out a second experiment, using a double-repeat within-subjects design.

## Methods

### Participants

27 healthy volunteers (18 women, mean age 21.3, SD age 2.59), with no history of neurological disorders, psychiatric disorders, or head injury were recruited from the local student population. Written informed consent was obtained from all volunteers. The study was approved by the University of Sussex local research ethics committee. Methods were carried out in accordance with the approved guidelines.

### Experimental design

The behavioural, data analysis and TMS components of each session were identical to that in experiment 1. However, unlike in experiment 1, the session was repeated for each participant 1 to 3 times on subsequent days, depending on performance on each day. The first day always involved bilateral cTBS to DLPFC (exactly like the DLPFC group in experiment 1). If the meta d’ – d’ score difference between pre- and post- cTBS administration on the first day was greater than 0.4 (in either direction, i.e. a metacognitive enhancement or impairment following DLPFC cTBS), then the participant was invited to a second day’s session, involving cTBS to the vertex. This threshold of 0.4 was the average effect found in the Rounis study. If in this second session the meta d’ – d’ score difference between pre- and post- cTBS administration was less than 0.2 (in other words, an appropriate control result) then the participant was invited to a 3^rd^ day’s session, where bilateral cTBS to the DLPFC was administered. If on this 3^rd^ day there was again a meta d’ – d’ score difference between pre- and post- cTBS administration greater than 0.4 then the participant was invited to a 4^th^ day’s session for cTBS to the vertex. In this way, we could rigorously explore the within subject likelihood of both a metacognitive impairment (or enhancement) following DLPFC cTBS *and* no metacognitive change following vertex cTBS, with a potential single subject replication of this pattern. metacognitive change following vertex cTBS, with a potential single subject replication of this pattern.

## Results

Of the 27 participants in this experiment, 9 were excluded because of extreme SDT values for type I or 2 HR and FAR (<0.05 or >0.95), and 1 further subject was excluded because of an exceptionally high type II FAR rate. The remaining 17 participants are summarized in table 5. Ten of these participants had no meta d’ changes on the first DLPFC session, and thus were not asked to return for subsequent sessions. Of the remaining 7 participants, 3 showed the expected impairment, while 4 showed a clear metacognitive enhancement following DLPFC cTBS. 6 of these 7 participants also showed a clear metacognitive change for the vertex control session, and thus were not asked to return for the 3^rd^ session (2^nd^ DLPFC). Only 1 participant that showed a clear DLPFC cTBS metacognitive change in the first session also showed no change for the 2^nd^ vertex cTBS session, and thus was brought back for the 3^rd^ DLPFC session. This session, unfortunately, included unstable SDT values, and thus the participant was not asked to return for a 4^th^ session. If these instabilities are ignored, though, the metacognitive change for the 3^rd^ session was very similar to the 1^st^ session. Both sessions, however, showed a robust *enhancement* of metacognition for this single subject following DLPFC cTBS, as opposed to the impairment found in the Rounis study.

**Table 5.** Experiment 2 values for meta d’ - d’ post TMS minus pre (above threshold results in bold)

| Sub no. | Session 1: DLPFC | Session 2: Vertex | Session 3: DLPFC | Session 4: Vertex |
| --- | --- | --- | --- | --- |
| 1 | **1.29** | **0.14** | **1.55 (Unstable)** |  |
| 2 | **1.00** | -0.80 |  |  |
| 3 | **-0.66** | 0.78 |  |  |
| 4 | **0.65** | 0.52 |  |  |
| 5 | **-0.61** | -0.46 |  |  |
| 6 | **0.40** | -0.42 |  |  |
| 7 | **-0.59** | -0.50 |  |  |
| 8 | 0.13 |  |  |  |
| 9 | -0.24 |  |  |  |
| 10 | -0.13 |  |  |  |
| 11 | -0.31 |  |  |  |
| 12 | -0.01 |  |  |  |
| 13 | 0.38 |  |  |  |
| 14 | 0.00 |  |  |  |
| 15 | 0.30 |  |  |  |
| 16 | -0.13 |  |  |  |
| 17 | -0.33 |  |  |  |

In summary, not a single subject of those 17 without instabilities showed both a metacognitive impairment in the DLPFC session, and no change in the vertex control session. The mean meta d’ – d’ change following DLPFC cTBS was 0.07, almost identical to that found in experiment 1 (0.06), and very different in magnitude and direction to that reported in the Rounis study (-0.35). When the first session DLPFC data from experiment 2 is combined with experiment 1, not only is there a clearly non-significant meta d’ – d’ difference between sessions (t(31)=0.14, p(1-tailed)=0.44; effect size Cohen’s d= 0.029), but also a Bayes Factor of 0.34, which is at the threshold for a robust null result (1/3).

## Discussion

We carried out two experiments to attempt to replicate Rounis and colleagues’ key finding that theta-burst TMS to DLPFC reduced metacognitive sensitivity [18]. We also attempted to extend these findings, by testing for a similar pattern of results for the PPC, and for only the left or right portion of the prefrontal-parietal network. In every case, we failed to demonstrate any modulatory effects of TMS on metacognition, when compared with an active TMS control site. No result even approached significance, on any of three measures of metacognition (type II d’, meta d’ and accuracy-confidence correlation), and all results either were close to, or passed a Bayes factor test for a robust confirmation of the null hypothesis. This was even the case when the control site was ignored, and the effects of DLPFC TMS were examined by themselves. We have therefore not only failed to replicate the Rounis result, but provided evidence from our own experiments that on this paradigm there is no modulatory effect of theta-burst TMS to DLPFC on metacognition.

There were several differences between our experiments and that of Rounis and colleagues. Perhaps the most notable divergence concerns data quality: we excluded subjects with unstable signal detection theory behavioural results (type I or 2 HR or FAR <0.05 or >0.95). Including such extreme results in the analysis is very likely to introduce instabilities in measures reliant on type I and II SDT quantities, including type II d’ and especially various implementations of meta-d’ (see [20] for a discussion of this issue). Specifically, since the z function (i.e. the inverse of the standard normal cumulative distribution) approaches plus or minus infinity as HR or FAR tends to 0 or 1, SDT measures such as meta d’ can take on extreme and highly inaccurate values with such inputs. In practice, we demonstrated from our data that unstable meta d’ – d’ values are significantly different from stable values, so that including them causes the sample to become non-Gaussian. Therefore, at the very least, any statistics on a sample including unstable SDT values should be non-parametric. Preferably, though, such data should be excluded entirely, to avoid false positive results. Indeed, when including these unstable values with parametric tests (as the Rounis study did), we did discover a significant effect of DLPFC TMS on meta d’, though one we know is invalid (namely a boost to metacognition in the control group).

Another difference between the current experiments and that of Rounis and colleagues is that we employed an active TMS control site, instead of sham TMS. Although our control results look similar to those of Rounis and colleagues, i.e. no modulatory effect of control TMS on metacognition, nevertheless it is still possible that this different approach to controls contributed to the different results. The DLPFC is amongst the most challenging sites to administer TMS, because of the peripheral facial nerves that can be activated, commonly causing facial twitching and minor pain. The participants in the Rounis study would have noticed a very dramatic difference between DLPFC and sham TMS, raising the possibility of demand characteristics influencing reported metacognitive deficit following DLPFC cTBS, compared to sham cTBS. The control paradigm used in the present study minimizes this potential psychological confound.

A third difference between our experiments and that of Rounis and colleagues is that they used relative visibility judgements for the metacontrast mask task, where participants attempted to give “clear” responses for 50% of their answers and “unclear” for the other 50%. Our experiments, instead, included non-relative responses of “random guess” and “at least some confidence” in the perceptual decision. We reasoned that this approach should have increased the sensitivity of our experiments to changes in metacognitive sensitivity, since our metacognitive labels are simpler for participants to categorise and process, with no working memory demand to maintain an equal number of answers for each label.

One intriguing positive finding from experiment 1 is that higher IQ participants tended to perform better on the objective part of the task, leading them to be presented with more difficult contrast levels. Although there were no similar relationships at the metacognitive level, this might be because higher IQ participants were effectively performing a more difficult perceptual task than lower IQ participants. The relationship between IQ and metacognition is still an open question. Nevertheless, future metacognitive studies in this area may benefit from recording IQ scores, or even restricting their sample to a narrow IQ range.

The result of Rounis and colleagues has recently gained a new significance given the emergence of so called “no-report” paradigms, which question the involvement of prefrontal-parietal regions in reportable perceptual transitions [36,37]. For instance, Frassle and colleagues used a binocular rivalry fMRI paradigm, and contrasted a standard report version with a ‘passive’ condition in which subjects did not explicitly report perceptual transitions, which instead were inferred from reflexive eye movements (nystagmus) [38]. In the passive condition, activity in the prefrontal parietal network was greatly reduced, especially in DLPFC, suggesting that many studies that associate this network with consciousness might be erroneously finding an association with the cognitive machinery necessary for overt response, rather than conscious perception per se. More recently, Brascamp and colleagues took this a step further, by using a binocular rivalry paradigm where reportability itself could be manipulated [36]. They used a clever stimulus arrangement which evoked perceptual transitions that were not perceived (and hence not reportable) by the subject: in other words, ‘change of perception’ without ‘perception of change’. In this condition, there were no detectable prefrontal parietal network changes at all, accompanying the perceptual transitions. Leaving aside the contentious issue of whether unreportable perceptual transitions should be classed as conscious, these recent studies are providing a fascinating alternative viewpoint to the previously dominant assumption that the prefrontal parietal network is critical for generating conscious contents. Our results are consistent with this emerging position.

However, there are alternative interpretations for our experiments. First, it may well be that cTBS of cortex, at the medically safe stimulation thresholds commonly employed (80% of active motor threshold) is just not intense enough to induce a subtle cognitive effect, such as a reduction in metacognitive sensitivity. To our knowledge, only a few published papers to date, besides that of Rounis and colleagues, have demonstrated the general efficacy of DLPFC cTBS in modulating cognitive performance [39-42]. First, Kaller and colleagues found only RT, rather than error effects on a planning task, when compared with a sham control [42]. Schicktanz and colleagues reported deficits on a 2 back working memory task, but – strangely - no deficits on a 0 or 3 back task, when comparing DLPFC cTBS with sham [41]. Neither of these studies, however, employed an active control, and so their effects could be attributed to participant expectations (i.e. demand characteristics). Ko and colleagues did use an active control in their study (vertex), but do not report behavioural comparisons between the DLPFC sites and the control site. Instead, they showed that the left DLPFC condition induced significantly more errors then the right DLPFC on the Montreal card sorting task [40]. Therefore, without a control comparison it is unclear whether they found a behavioural deficit at all. Finally, Rahnev recently applied cTBS to a range of sites on a metacognitive task and found that cTBS actually boosted metacognition for DLPFC and anterior prefrontal cortex, compared to a control site [39]. In other words, these results run counter to those of Rounis and colleagues.

Adding other tasks associated with the prefrontal parietal network to metacognitive paradigms like ours, for instance involving working memory, may therefore be useful. If we had found clear working memory impairments following DLPFC cTBS, for instance, *but not metacognitive impairments*, this would have demonstrated the general effectiveness of DLPFC cTBS. Given that we were focusing on closely replicating the Rounis paradigm, we were unable to include these extra conditions, but future experiments that further investigate these effects may consider modifying the paradigm in this way.

A second alternative interpretation for our null result is that cTBS of cortex, especially when it involves highly flexible, semi-redundant areas like the prefrontal parietal network, might, after all, induce rapid functional and/or structural plasticity effects that compensate for any possible functional impairment. For instance, when cTBS was applied bilaterally to DLPFC in the current experiments, it may be that posterior parietal cortex transiently takes on a larger role in metacognitive decisions while DLPFC neurons were being moderately suppressed.

Finally, we recognize that our study may have differed from Rounis et al. in how effectively the DLPFC was targeted by TMS stimulation. Factors affecting targeting efficacy include variations in measurement of stimulation site, coil location and orientation, head shape, and the like. Although we assumed, given we used exactly the same TMS targeting method as Rounis and colleagues, that such variability would have been roughly similar between studies, future studies may partially address these issues by using individual structural MRI data to guide TMS stimulation in combination with ‘neuronavigation’ methods that allow targeting of TMS to specific cortical regions with increased fidelity [43]. However, the fact that we did not observe metacognitive impairment reliably in any single subject in experiment two speaks against interpreting our null results simply in terms of missing the DLPFC during cTBS, since at least some of these subjects should have had cTBS closely over DLPFC (site locations for each condition were fixed between sessions).

Although it is difficult to know which of these interpretations is more likely, our results nevertheless indicate that the cTBS approach is not, so far, sensitive enough to establish a causal link between DLPFC and metacognitive processes. They also emphasize the importance of giving careful methodological consideration both to the design of effective control conditions, and (especially for metacognitive studies), of excluding unstable data which may otherwise confound sophisticated statistical analyses. Overall, our results contribute to the evolving discussion concerning the role of the prefrontal-parietal network in conscious visual perception. Future studies that take into account both our data and the Rounis et al results, alongside emerging “no-report” paradigms, may yet resolve this critical issue in consciousness science and metacognition.

## Acknowledgements

We thank the Alex Henderson and Arin Baboumian for all their work collecting the data, Ryan Scott and Zoltan Dienes for helpful theoretical discussions, and Justyna Hobot on many constructive comments on an earlier draft. This work was supported by The Dr Mortimer and Theresa Sackler Foundation. ABB is funded by EPSRC grant EP/L005131/1.

## Author Contributions

DB devised the studies, DB and DS carried out the studies and analysed the data, DB, AB and AKS contributed statistical and theoretical points, and DB, DS and AKS wrote the paper.

